# *Ide* copy number variant does not influence lesion size and mortality in two C57BL/6J mouse models of cerebrovascular ischemia nor in human cerebrovascular disease. An exploratory study

**DOI:** 10.1101/2024.05.15.593342

**Authors:** Marco Foddis, Sonja Blumenau, Susanne Mueller, Clemens Messerschmidt, Clarissa Rocca, Alistair T Pagnamenta, Katarzyna Winek, Matthias Endres, Andreas Meisel, Arianna Tucci, Jose Bras, Rita Guerreiro, Dieter Beule, Ulrich Dirnagl, Celeste Sassi

## Abstract

Contrary to the common belief, the most commonly used laboratory mouse inbred strains are shaped by a distinctive genetic and phenotypic diversity. In the past 10 years next generation sequencing unveiled a wide spectrum of genetic variants in different mouse inbred strains and the meticulous observation of researchers pointed to a variegate intra-and inter-strain phenotypic diversity. Although a genotype-phenotype correlation has been described for some traits, the relationship between several endophenotypes and causative genetic variability remains still unknown. Recently, we characterized the brain collateral plasticity in two brain ischemia C57BL/6J mouse models (i.e bilateral common carotid artery stenosis [BCCAS] and 60-min transient unilateral middle cerebral artery occlusion [MCAO]) and observed a Mendelian-like fashion of inheritance of the posterior communicating artery (PcomA) plasticity. Interestingly, a copy number variant (CNV) spanning *Ide* locus was reported to segregate in an analogous Mendelian-like pattern in the C57BL/6J colonies of the Jackson Laboratory. Given the critical role of *Ide* in vascular plasticity, *Ide* CNV was an excellent candidate to explain PcomA variability in C57BL/6J inbred mice. To investigate this hypothesis, we applied a combination of complementary techniques (i.e T2-weighted magnetic resonance imaging [MRI], time of flight [TOF] angiography [MRA], cerebral blood flow [CBF] imaging and histology) to characterize the collaterome in C57BL/6J BCCAS and MCAO mice and performed on these Taqman genotyping, exome sequencing, and RNA sequencing. We report an *Ide* CNV in a BCCAS mouse with 2 patent PcomAs. We then investigated the hypothesis that *IDE* gain and loss of function mutations may have influenced the vascular phenotype in a cohort of 438,250 cases and controls (UK Biobank) and 15,790 neurological patients (Genomics England), respectively. We identified four *IDE* CNVs resulting in a loss of function (LoF) in one patient with hereditary ataxia, a patient with hereditary congenital heart disease and two healthy individuals. In addition, we report four *IDE* LoF point mutations (p.Leu5X, p.Met394ValfsX29, p.Pro14SerfsX26, p.Leu889X) present in controls or inherited from healthy parents. *Ide* CNV and LoF variants do not crucially influence PcomA variability in C57BL/6J inbred mice and do not cause a vascular phenotype in humans.

## INTRODUCTION

Despite seven decades of inbreeding through several hundreds of brother-sister mating generations, inbred mice widely used as experimental model of disease remain only virtually and utopically isogenic ^1, 2, 3, 4, 5^. In the past ten years deep genotyping and next generation sequencing triggered a turbulent wave of genetic discoveries and unveiled a wide range of intra-and interstrain genetic differences: from synonymous non-coding variants to kilo-megabase copy number variants ^1, 5^. This demonstrates that inbred mice, contrary to the common belief, should be considered as members of the same extended multigenerational family, rather than homozygotic twins. In support of these genetic studies, a colourful array of phenotypic traits has been described: from macroscopic differences such as the hair colour, body size, density of the bone mass, to metabolic, behavioural and functional phenotypes, that can only be observed under specific circumstances ^6, 3, 7, 2^. Thus, implying that during decades traits have been positively selected and evolutionary forces gave rise to diverse substrains. Although critical genetic factors have been already found for few of these endophenotypes ^3, 8^, for several of these the genotype-phenotype correlation is yet to be discovered. Among these, we recently reported the variability of the posterior communicating artery (PcomA) patency during acute and subacute brain hypoperfusion in 2 brain ischemia mouse models (bilateral common carotid artery stenosis [BCCAS] and middle cerebral artery occlusion [MCAO]) ^9^. Importantly, the PcomA recruitment is a very dynamic process, represents the main variable survival mechanism and the main determinant of stroke lesion volume and recovery in both models and, in line with previous studies, segregated within the C57BL/6J strain in a Mendelian-like fashion (70% of the mice displayed 1 prominent PcomA, 30% no PcomA and 20% presented 2 very prominent PcomAs) ^9, 10^. Interestingly, among the genetic differences reported in the C57BL/6J strain colonies from the Jackson Laboratory, a ∼ 112 Kb CNV on chromosome 19, encompassing the insulin degrading enzyme gene (*Ide*), associated to a significantly increased *Ide* expression has been analogously reported to be inherited within the C57BL/6J strain in a Mendelian-like pattern (64% of mice heterozygous for the CNV, 23% without CNV and 13% homozygous for the CNV) ^1^. Considering that CNVs represent the main mechanism of genome evolution ^11,12^, given *Ide* expression in brain vessel endothelial cells ^13^, *Ide* CNV absence in Balb mice ^14, 1^, which are characterized by a poor collateralization during acute brain hypoperfusion compared to the C57BL/6J strain^15, 16^ and the overlapping pattern of segregation of *Ide* CNV and PcomA patency ^9, 10^ within C57BL/6J strain, we hypothesize that CNV in *Ide* may explain the diversity of PcomA calibre in the same strain.

To investigate this hypothesis, we applied a combination of complementary techniques (T2-weighted magnetic resonance imaging [MRI], time of flight [TOF] angiography [MRA], cerebral blood flow [CBF] imaging and histology) to characterize brain arterial collaterals in C57BL/6J BCCAS and MCAO mice and genetically characterized these performing Taqman genotyping, exome sequencing and RNA sequencing (**Figure 1**). We then investigated *IDE* gain and LoF in cohorts of neurological patients and controls (454,040 individuals from the UK Biobank and Genomics England).

**Figure 1.**
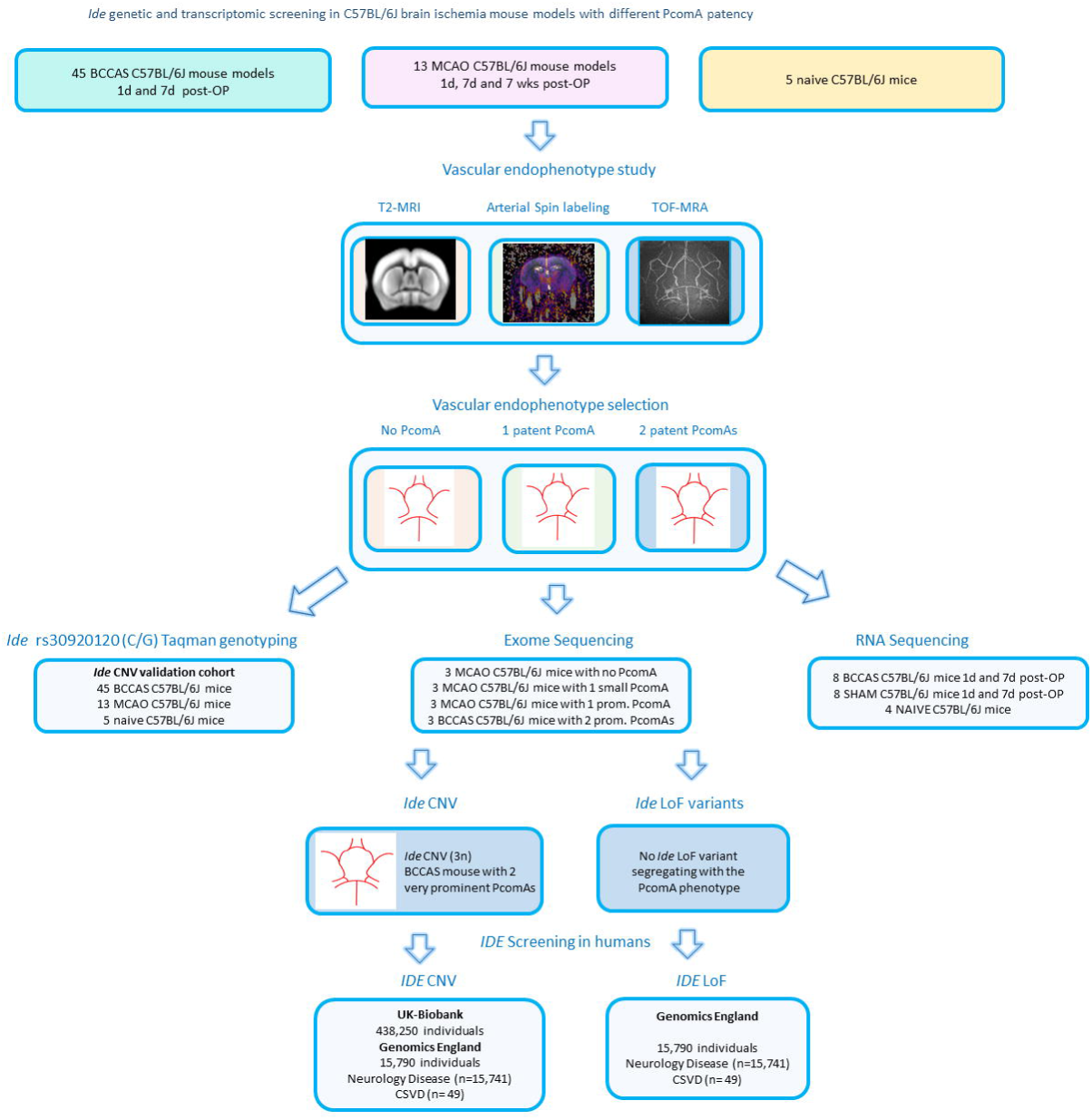
Pipeline followed in our study. We used 2 acute ischemia brain mouse models (BCCAS [bilateral common carotid artery stenosis] and MCAO [middle cerebral artery occlusion]) and studied the posterior communicating artery phenotype (PcomA) using T2 weighted MRI, cerebral blood flow measurement through arterial spin labeling and TOF MRA. We then selected BCCAS and MCAO mice with different PcomA phenotype for *Ide* rs30920120 Taqman genotyping, exome sequencing and RNA sequencing. We identified one BCCAS mouse with 2 prominent PcomAs carrying a *Ide* CNV. Although this *Ide* CNV did not segregate with PcomA patency phenotype, we screened also the *IDE* gain and loss of function mutations in a human cohort of cases and controls from 2 databases (UK-Brain Bank and Genomics England), where *IDE* CNV and loss of function mutations are very rare and not likely to play a critical influence on vascular phenotypes.

Importantly, linking the brain arterial collateral plasticity to specific genetic variants has the potential to provide a unique window into the more complex genetic-phenotypic variability of humans.

## MATERIALS AND METHODS

### Animals, experimental design and exclusion criteria

Experiments were approved by the Landesamt für Gesundheit und Soziales and conducted according to the German Animal Welfare Act and institutional guidelines. 54 and 18 male C57/BL6 J mice (purchased at 8 weeks of age, Charles River, Germany and 10 weeks of age Janvier France, respectively) were housed in a temperature (22±2°C), humidity (55±10%), and light (12/12-hour light/dark cycle) controlled environment. The animals are subject to brain hypoperfusion between 9 and 13 weeks of age (n=50, BCCAS= 35, MCAO= 15) or were used as controls (naïve=11; BCCAS sham= 8; MCAO sham=3).

The only exclusion criterion was death during MRI due to wrong placement of the animal in the scanner and led to exclusion of 2 MCAO and 2 sham animals (MCAO group), resulting in final analyzed sample size of MCAO = 13 and MCAO sham =1.

BCCAS mice were imaged before surgery, 24 hours and 1 week post-surgery and imaging for 7 weeks were re-analyzed from a previous study ^17^. MCAO mice were imaged 24 hours, 1 week, 4 weeks and 7 weeks post-surgery for angiography and estimation of cerebral blood flow (CBF) using arterial spin labeling. At 2 days and 8 days (BCCAS) and 7 weeks (MCAO) tissue was processed for immunohistochemistry

### RNA Isolation for mouse samples and Real-Time PCR

Total RNA from C57BL/6J BCCAS and MCAO blood was isolated using QIAamp RNA Blood Mini Kit (QIAGEN). The quality and the concentration of the total RNA was determined using a Nanodrop Spectrometer (A260:A280 and A260:A230 ratios). For real-time PCR analysis, 750ng total RNA from each sample was used for first-strand cDNA synthesis using SuperScript III (Invitrogen). cDNA from each sample was amplified via real-time PCR and normalized against Actin, using LightCycler 480 Instrument II (Roche). mRNA levels for each experimental group were quantified using the comparative CT method.

### RNA sequencing

Eight BCCAS, 8 sham and 4 näive mice were sacrificed with cervical dislocation 2 days and 8 days post coil insertion surgery, followed immediately by postmortem dissection of the prefrontal cortex, striatum and hippocampus from one hemisphere. The other hemisphere was preserved for immunohistochemistry. The dissected tissues were immersed in RNA later and stored at −80°C for later use for mRNA-Sequencing. Total RNA was extracted using miRNeasy Kit (Qiagen, Cat # 217004). Total RNA quality was assessed with the use of Bioanalyzer. Average RIN (RNA Integrity Number) of our samples was 9. Next Generation Sequencing mRNA libraries were prepped with Illumina TruSeq RNA Library Preparation Kit (Illumina, Cat # RS-122-2001).

### Mouse cohort for exome sequencing

Our mouse cohort was composed of 9 MCAO mice and 3 BCCAS mice with different collateral circulation phenotype (**Table 1**, **Figure 1**).

**Table 1.**
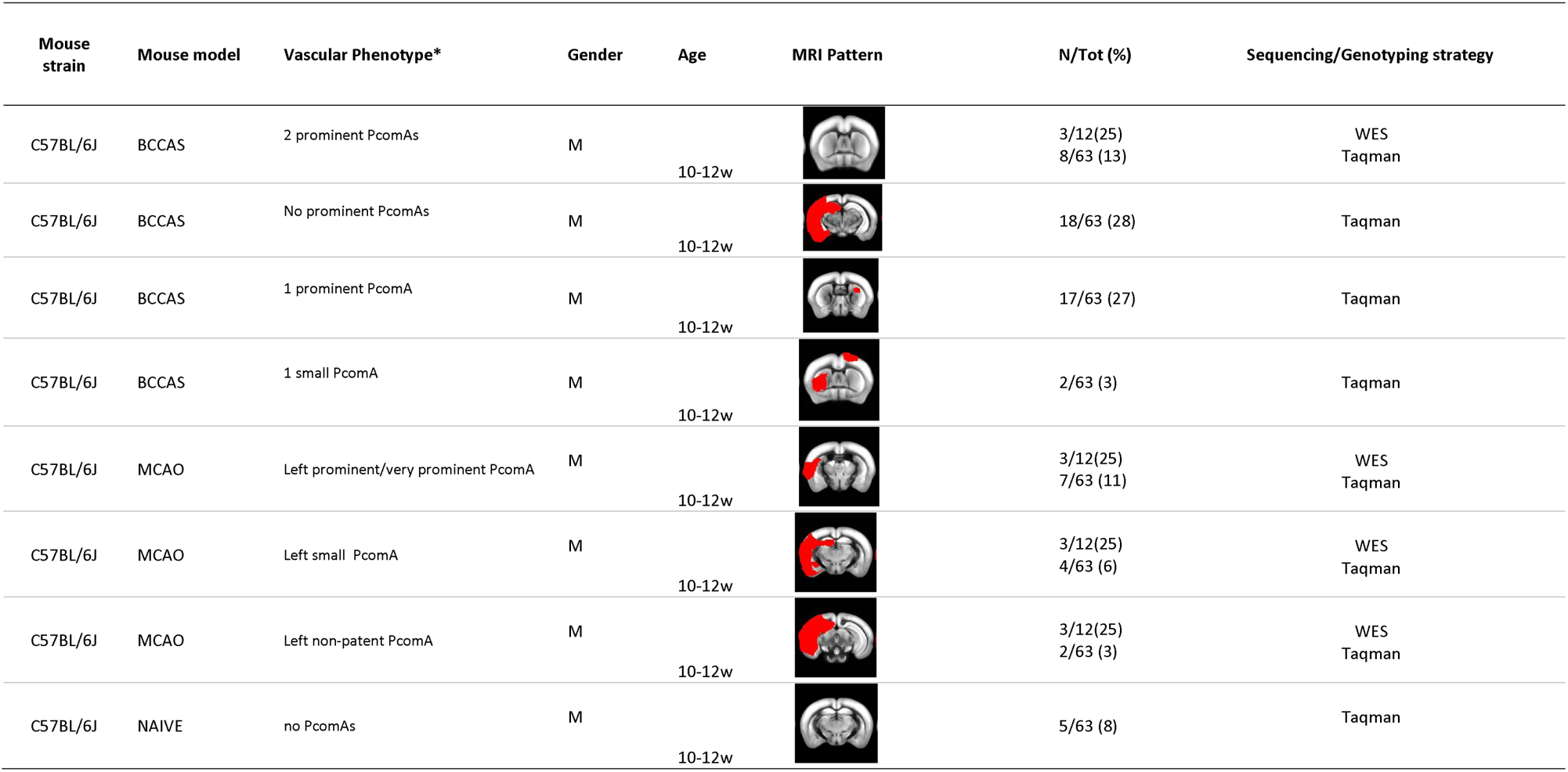
Exome sequencing and *Ide* rs30920120 (C/G) Taqman genotyping cohort description. BCCAS, bilateral common carotid artery stenosis; MCAO, middle cerebral artery occlusion; PcomA, posterior communicating artery; w, weeks; WES, whole exome sequencing; N,number. PcomA classification is based on the PcomA/Basilar Artery diameter ratio ^1^, already described.

| Mouse strain | Mouse model | Vascular Phenotype* | Gender | Age | MRI Pattern | N/Tot (%) | Sequencing/Genotyping strategy |
| --- | --- | --- | --- | --- | --- | --- | --- |
| C57BL/6J | BCCAS | 2 prominent PcomAs | M | 10-12w |  | 3/12(25)<br>8/63 (13) | WES<br>Taqman |
| C57BL/6J | BCCAS | No prominent PcomAs | M | 10-12w |  | 18/63 (28) | Taqman |
| C57BL/6J | BCCAS | 1 prominent PcomA | M | 10-12w |  | 17/63 (27) | Taqman |
| C57BL/6J | BCCAS | 1 small PcomA | M | 10-12w |  | 2/63 (3) | Taqman |
| C57BL/6J | MCAO | Left prominent/very prominent PcomA | M | 10-12w |  | 3/12(25)<br>7/63 (11) | WES<br>Taqman |
| C57BL/6J | MCAO | Left small PcomA | M | 10-12w |  | 3/12(25)<br>4/63 (6) | WES<br>Taqman |
| C57BL/6J | MCAO | Left non-patent PcomA | M | 10-12w |  | 3/12(25)<br>2/63 (3) | WES<br>Taqman |
| C57BL/6J | NAIVE | no PcomAs | M | 10-12w |  | 5/63 (8) | Taqman |

The study of the PcomA role during acute hypoperfusion followed Martin et al. PcomA classification ^18, 19^ and has been already described ^9^. Briefly, this identifies 4 PcomA classes, based on the ratio between PcomA and basilar artery (BA) diameter: 1) PcomA <10% of BA; 2) PcomA 11-20% of BA; 3) PcomA 21-30% of BA and 4) PcomA >30% of BA. We identify class 1 and class 2 as ‘non-patent’, class 3 as ‘small ‘, class 4 as ‘prominent’ and included a fifth class, represented by PcomA>60% of BA, described as ‘very prominent’.

The diameters of the PcomAs were measured at the smallest point and the diameter of the BA was measured proximal to the superior cerebellar arteries both for the Evans Blue and fluorescent WGA stainings (MCAO mice) or only with for Evans Blue staining (BCCAS mice) with ImageJ. The diameter of the PcomAs as a percentage of the diameter of the BA was calculated and used in the analysis as previously described ^18^ ^19^.

In our mouse cohort, 3 BCCAS mice with 2 very prominent PcomAs, together with 9 MCAO mice, characterized by different left PcomA calibre: a) 3 MCAO mice with prominent-very prominent left PcomA, that developed small ischemic lesions (≈5-10% of the left hemisphere), mostly affecting ventral areas (prefrontal cortex, striatum and ventral hippocampus), and were characterized by the most favourable stroke outcomes (**Table 1**); b) 3 MCAO mice with non-patent PcomA, which died few hours post surgery and c) 3 MCAO mice with small PcomA, that survived the surgery but developed monolateral severe left strokes affecting up to one third of the left hemisphere and including also dorsal areas (orbital cortex and cerebellum) (**Table1**) were selected for our exome sequencing study.

Given the extreme inbreeding of the C57BL/6J strain, carefully inbred for over seventy years through more than 200 generations of brother-sister mating ^1^, and the likely minimal influence of environmental factors, these mice were genetically considered as members of the same large multigenerational family coming from a small and isolated village. Moreover, the selection of extreme phenotypes (absent-small PcomA vs prominent-very prominent PcomA), allowed us to reach an high power for the detection of rare variants with large effect size, despite the small sample size ^20^ ^21^, although no formal sample size/power calculation was performed due to the exploratory nature of the study.

### Exome sequencing in C57BL/6J mice

We performed whole exome sequencing (WES) on a cohort of 12 C57BL/6J mice (9 MCAO and 3 BCCAS). DNA was extracted from cerebellum using standard protocols. Library preparation for next generation sequencing used 50 ng DNA. Exome libraries were prepared using Nextera® Rapid Capture Exome and Kit (4 rxn × 12 plex, FC-140-1002) and Nextera DNA Library Prep Kit (FC-121-1030). The DNA library was then hybridized to an exome capture library (Nextera, Illumina Inc.) and precipitated using streptavidin-coated magnetic beads (Nextera, Illumina). Exome-enriched libraries were PCR-amplified, and then DNA hybridized to paired-end flow cells using a cBot (Illumina, Inc.) cluster generation system.

The WES libraries were sequenced paired-end 75 bp on Illumina HiSeq 4000 with a median of 60.5 million reads per library.

### Bioinformatics RNA sequencing in C57BL/6J mice

Processing, quality assessment and analysis of RNAseq data was carried out using a custom pipeline. We aligned paired end reads with STAR ^22^ against the GRCm38.p4 genome using gencode.vM12 annotation ^23^ (http://www.gencodegenes.org/mouse_releases/12.html), excluding alternative scaffolds and patches. Gene counts were determined using HTSeq ^24^. Testing for differential gene expression and cerebral blood flow and gene-expression correlation was done using DESeq2 ^25^. Genes were counted as differentially expressed where they had a moderated fold change of 2 or more, contrasting coil to shame samples and where their false discovery rate (FDR) adjusted p-value was below 0.05.

### Bioinformatics, exome sequencing in C57BL/6J mice

The reads were aligned using BWA-MEM v0.7.15 ^27^ to the reference GRCh37 (hs37d5.fa), separate read groups were assigned for all reads from one lane, and duplicates were masked using Samblaster v0.1.24^28^. Standard QC was performed using FastQC (http://www.bioinformatics.babraham.ac.uk/projects/fastqc). The variants were then called using GATK UnifiedGenotyper v3.7^29^ and annotated using Jannovar v0.24^30^ using RefSeq v105 exons.

For the CNV analysis of the WES data, Cnvkit (Talevich et al., 2014) in batch mode was used in a matched fashion as described in their manual for WES data.

All methods were carried out in accordance with relevant guidelines and regulations.

### *IDE* genetic screening in a human cohort

#### CNVs overlapping IDE in the UK Biobank and in the Genomics England databank

CEL files from 438,250 individuals were downloaded and CNVs were called using both Affymetrix Power Tools and PennCNV. These individuals were not selected based on any phenotype or diagnosis.

CNVs were considered if including at least 10 SNPs and if at least 50kb in length. Adjacent CNVs were merged based on the default parameters, and CNVs were excluded if they were overlapping telomeres, centromeres, known segmental duplications, immunoglobulin, or T cell receptor loci. PennCNV was used to identify genes which overlapped the CNV calls.

After QC, a total of 7 individuals were identified as potential CNV carriers overlapping *IDE*. To evaluate the authenticity of these calls, B-allele frequency and Log R Ratios were used to plot the CNVs. Plots were then generated covering IDE +/−5Mb.

Copy number variants in IDE gene in the genomics England database were investigated as previously described ^31^

#### Loss of function variants in the Genomics England database

We analysed data from the 100,000 Genomes Project (Genomics England, The National Genomics Research Library v5.1, Genomics England. doi:10.6084/m9.figshare.4530893/7. 2020.) for families where affected individuals harboured loss of function variants in *IDE*. All genomes from probands and affected family members (n=15,790) recruited under the Neurology Disease group (n=15,741) and Familial cerebral small vessel disease (n= 49) in the 100KGP were annotated (NM_004969.4) and analysed for *IDE* variants. Then we filtered the dataset for loss of function variants with allele frequency below 0.01

### MRI Data Analysis in C57BL/6J mice

Cerebral blood flow and angiography CBF maps were calculated using the Perfusion ASL macro in Paravision 5.1 software via the T1 method using a blood T1 value of 2100 ms and a brain blood partition coefficient of 0.89 mL/g 1,2. A custom written Matlab Release 2013a (MathWorks, Natick, MA, USA) script extracted the CBF maps from Paravision, and prompted manual delineation of regions of interest (ROI) in the striatum and prefrontal cortex. The resulting CBF values were expressed in mL/min/100g.

Angiography images were analyzed as previously described ^17^. Briefly, the spatial dimensions or the raw data was up-scaled by a factor of ten, exported into FSL software (Analysis Group, FMRIB, Oxford, UK), and the FLIRT tool was used for coregistration. Registered images were exported into ImageJ freeware (National Institutes of Health) and a maximum intensity projection (MIP) of the Circle of Willis was prepared with a custom plugin. A threshold (14 000 in 16 bit images, i.e. ∼43% of max) was used to create a binary image of the MIP so that the number of voxels in the Circle of Willis could be counted and expressed in µm2.

Ventricle to brain ratio (VBR) and hippocampal size were calculated from the T2 weighted images. Outlines of all structures were manually delineated on a slice by slice basis in ImageJ, and total volumes of each were calculated by multiplying each area by slice thickness (0.50 mm) and summation.

Fisher’s exact test on lesion volume and CBF values was performed. A p-value of 0.05 was set as a nominal significance threshold. All computations, were performed in R (version x64 3.0.2, http://www.r-project.org/).

### Taqman genotyping in C57BL/6J mice

*Ide* genotype rs30920120 (C/G) was assayed using LightCycler 480 Instrument II (Roche) or the TaqMan method (Applied Biosystems Inc. [ABI], Foster City, CA,USA). SNP-specific primers and probes were designed by Thermofischer or ABI (TaqMan genotyping assays). TaqMan real-time polymerase chain reaction assays (PCR) consisted in 2.5 ul of Fast Master Mix (Roche), 0.125 ul of assay, 0.375 ul of water and 2ul of DNA at 5ng/ul. The 5μl total volume reaction was loaded in 384-well plates and was performed in a LightCycler 480 Instrument II (Roche), using a cycling program of: 95°C for 10 min; 40 cycles of 95°C for 15 sec and 60°C for 1 min. Six positive controls, one for each genotype, and one negative control (water) were included in each plate and were consistently called correctly.

### Methods to prevent bias

This is an exploratory, descriptive study. Sample sizes were not based on *a priori* power calculation. Mice were randomized to receive hypoperfusion. The study was only partially blinded.

## RESULTS

In this study we tested the hypothesis that the phenotypic correlate of *Ide* duplication and gene expression variability reported in C57BL/6J inbred strain from the Jackson Laboratory ^1^ may have been the different dynamic PcomA patency degree, described in the same strain with overlapping frequency^9^.

To test this hypothesis we used a combination of complementary genetic techniques (Taqman genotyping, exome sequencing and RNA sequencing) in two different C57BL/6J mouse models of brain acute and subacute hypoperfusion: BCCAS and MCAO (**Figure 1**).

### Taqman genotyping of Ide rs30920120 of BCCAS and MCAO and naive C57BL/6J mice with different PcomA patency features. Validation cohort

We selected the *Ide* rs30920120 probe, corresponding to the coding SNP C/G on chromosome 19 at *Ide* Locus, and described by Watkins-Chow and colleagues as not showing identical heterozygosity in the C57BL/6J mouse strain instead falling in 2 discrete genotype classes differing in their signal intensity ratio ^1^. We performed Taqman genotyping using the probe rs30920120 on a cohort of 63 C57BL/6J mice: 45 BCCAO mice (69%) 1 day or 7 days post surgery, 13 MCAO mice (23%) 1 day, 7 days or 7 weeks post-surgery and 5 naïve mice (8%), which did not display any patent PcomA (**Table 1**). All the samples displayed identical heterozygosity and overlapping signal intensity, without falling into two discrete genotype classes, as described in Jackson laboratory C57BL/6J mice ^1^(**Figure 2, Table S1**).

**Figure 2.**
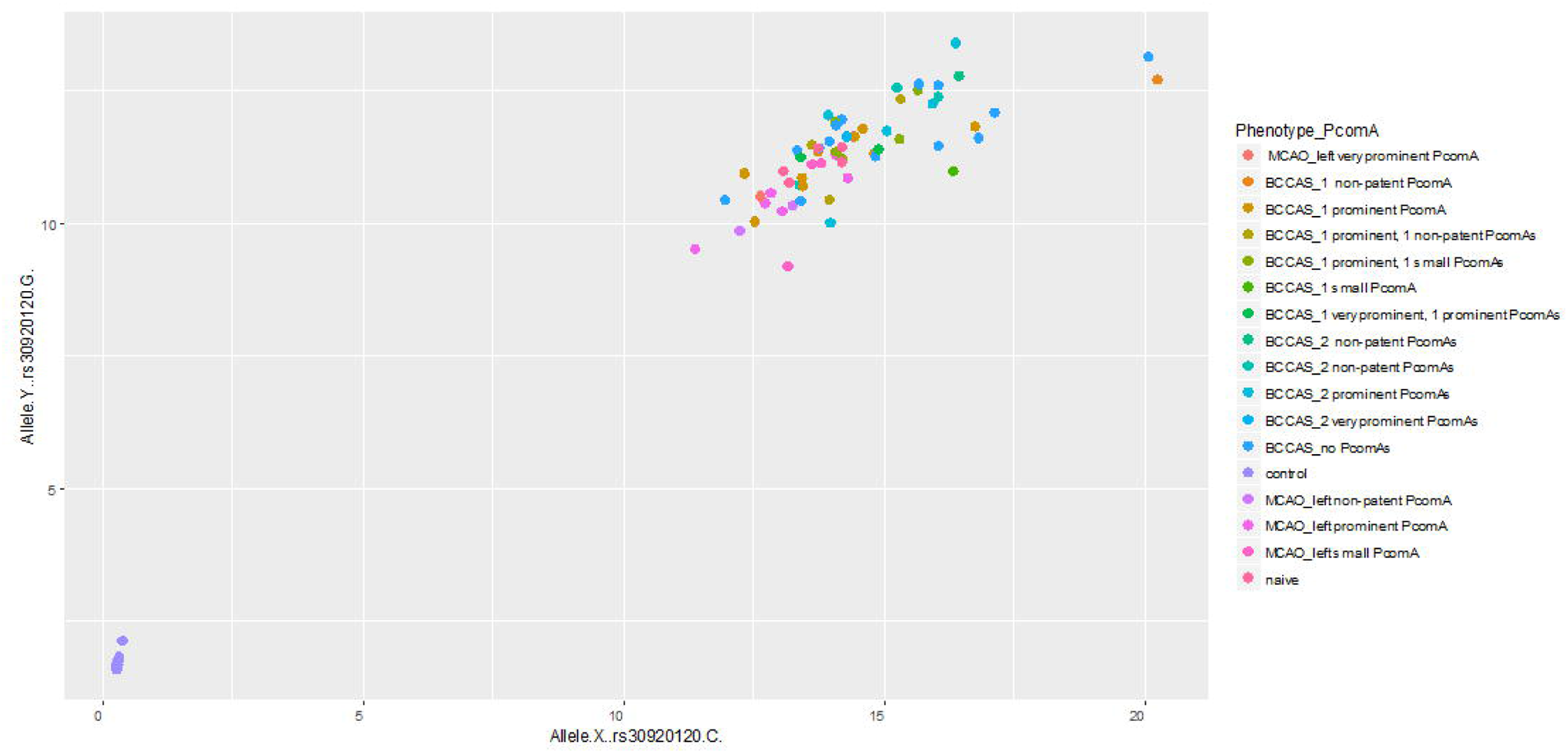
*Ide* rs30920120 (C/G) Taqman genotyping in a cohort of BCCAS and MCAO and naive mice with different PcomA phenotype.

We then performed whole exome sequencing in 12 C57BL/6J MCAO and BCCAS mice that showed phenotypically significantly different stroke lesion sizes, collateral blood flow and arterial brain collateral recruitment pattern.

### Ide CNV detection in exomes of BCCAS and MCAO mice with different PcomA phenotype

We performed exome sequencing in 9 MCAO and 3 BCCAS mice with diverse PcomA caliber and investigated the hypothesis that PcomA spectrum size, ranging from no PcomA/non-patent PcomA to very prominent PcomAs may have been determined by coding missense mutations in *Ide* (**Figure 1**, **Table 1**).

We report an *Ide* CNV (3n) in a BBCAS mouse with 2 effective PcomAs (**Figure 3**) and not segregating with the patent-PcomA phenotype.

**Figure 3.**
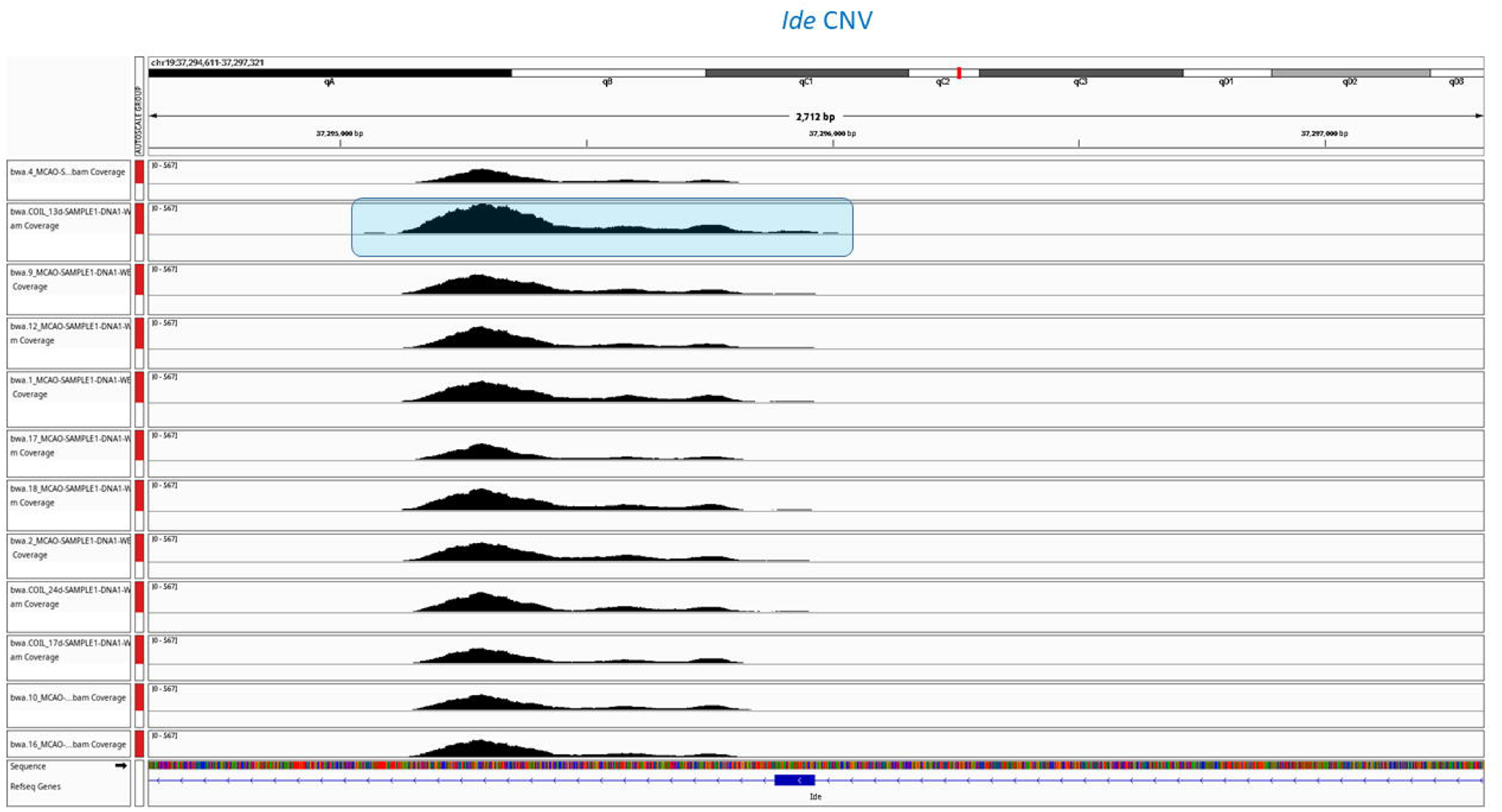
Copy number variant analysis of *Ide* Locus, based on exome sequencing data on 12 BCCAS and MCAO mice with different PcomA phenotype.

We next focused on loss of function mutations in *Ide* as another likely genetic mechanism potentially explaining intra-strain vascular differences in our mouse exome cohort and did not detect any *Ide* LoF mutation segregating with the PcomA phenotype.

Finally, since the *Ide* CNV reported in the Jackson Laboratory colonies corresponds to a proportional increase in *Ide* gene expression we further examined firstly, whether during brain hypoperfusion there was a significant increase of *Ide* expression in BCCAS mice in the most hypoperfused brain areas and secondly, whether there may have been a correlation between *Ide* differential gene expression, ischemic lesion sizes and cerebral blood flow recovery (**Figure 4-5**). Therefore we performed RNA sequencing during acute (1d) and subacute (7d) brain hypoperfusion in hippocampus, striatum and prefrontal cortex of C57BL/6J BCCAS mice with different stroke lesion sizes, cerebral blood flow recovery and therefore PcomA patency (**Figure 1**).

**Figure 4.**
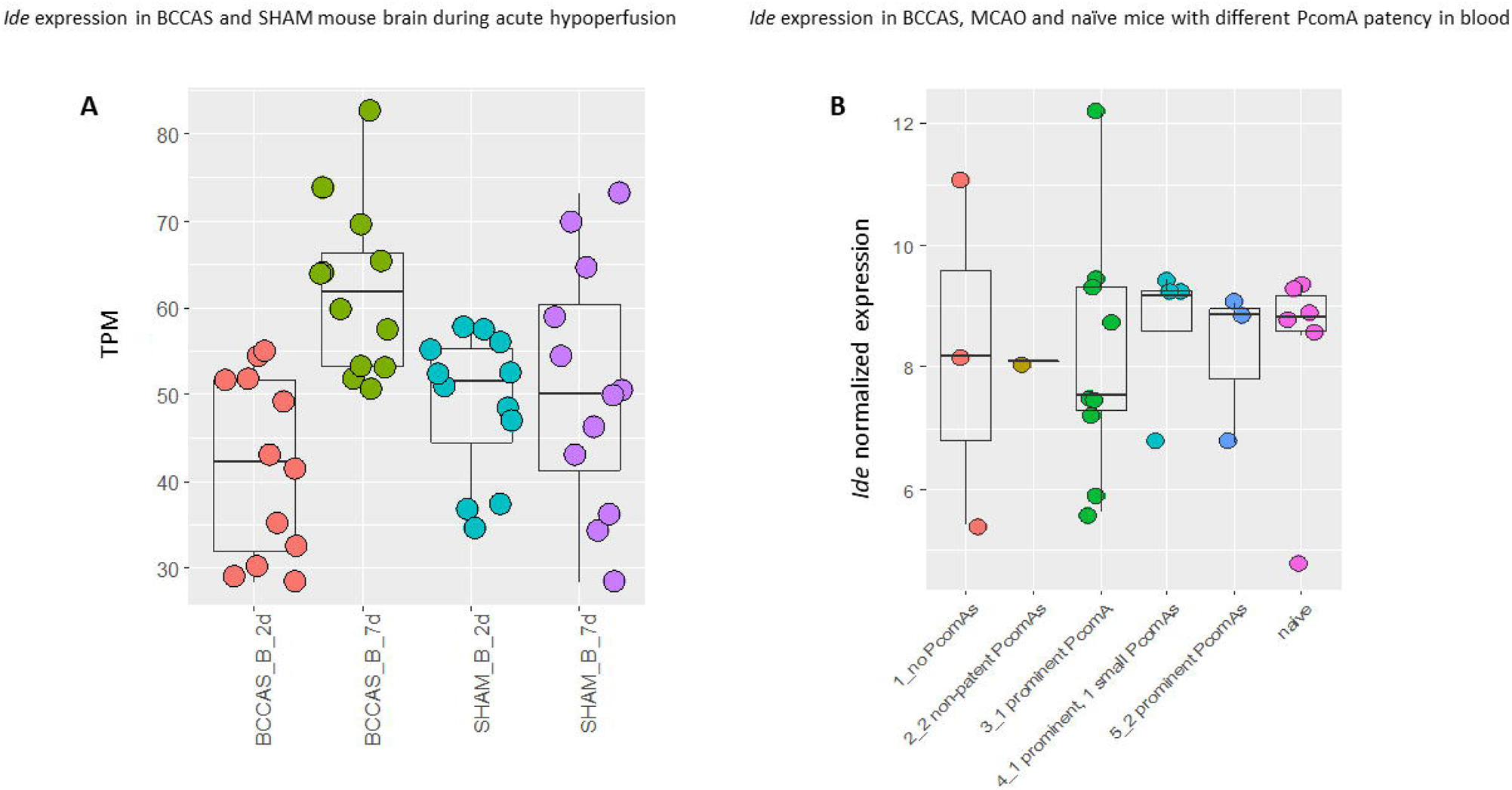
**A.** RNA sequencing *Ide* brain expression analysis in BCCAS and SHAM mouse brain (prefrontal cortex, hippocampus and striatum) during acute hypoperfusion. **B.** Real-TIME PCR *Ide* expression analysis in BCCAS, MCAO and naïve mice with different PcomA patency in blood during acute hypoperfusion.

**Figure 5.**
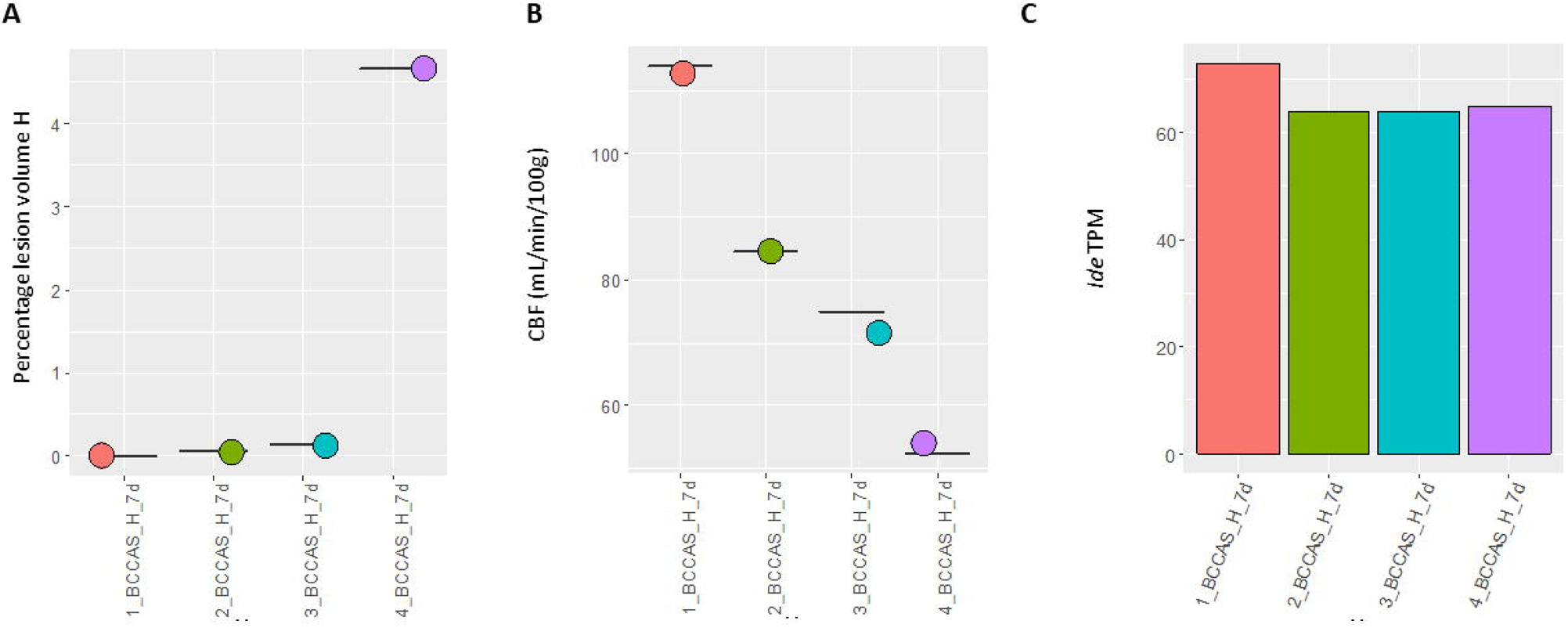
*Ide* hippocampal gene expression analysis based on RNA sequencing data in BCCAS mice during subacute hypoperfusion (7d) with different hippocampal lesion sizes and cerebral blood flow recovery. **A.** Percentage of hippocampal lesion volume in BCCAS mice 7 days post-surgery. **B**. CBF measurements in BCCAS mice in hippocampus 7 days post-surgery (CBF, ml/min/100g). **C**. *Ide* gene expression in the hippocampus of BCCAS mice 7 days post-surgery.

### Ide expression in hippocampus, striatum and prefrontalcortex in BCCAS C57BL/6J mice during acute and subacute brain hypoperfusion

We did not detect a significant *Ide* differential expression during acute and subacute brain hypoperfusion in hippocampus, striatum and prefrontal cortex between BCCAS and SHAM or naïve mice as well as in blood samples of BCCAS, MCAO naïve mice with different PcomA patency phenotype (|Fold change|<2, FDR p-value>0.05) (**Figure 4 A-B**). Moreover, we do not report any correlation between *Ide* expression, severity of the stroke lesion size and rapidity of cerebral blood flow recovery (**Figure 5 A-C**). Thus, implying that brain diverse collateralization is not shaped by *Ide* expression.

### CNVs overlapping IDE in the Genomics England database and UK Biobank

We found 4 *IDE* CNVs in the Genomics England database: two *IDE* CNVs (Chr10:-92556744-92561449, Chr10:92557209-92565408) were detected in one patient with hereditary ataxia and with syndromic congenital heart disease, respectively and two *IDE* CNVs (Chr10:-92556744-92561449, Chr10:92557209-92565408) were found in two healthy controls (**Table 2**). All of these *IDE* CNVs were identified in a nonsense-mediated mRNA transcript (NM_001322795), expressed at very low levels in the skin (Gtex data), the tissue where IDE is most abundant.

**Table 2.** *IDE* LoF and *IDE* CNV mutations detected in the 15.790 genomes from the 100,000 Genomes Project (Genomics England). Aa, ammino-acid; Gen, genotype; Het, heterozygous.*Also present in healthy controls

| Position | rsID | Ref/Alt | cDNA | Aa | Gen | gnomAD_AF | ClinVar | GenE (%) | Phenotype |
| --- | --- | --- | --- | --- | --- | --- | --- | --- | --- |
| chr10:92574007 | rs533083105 | AG/A | c.13del | p.Leu5X | Het | 0,000958 | Not present | 41/15,790 (0.2) | Neurology and Neurodevelopmental disorders/Cardiovascular Disorders* |
| chr10:92507639-92507640 | rs749353444 | CAT/C | c.1180_1181del | p.Met394ValfsX29 | Het | 0,00005172 | Not present | 1/15,790 (0) | Neurology and Neurodevelopmental disorders* |
| chr10:92573971-92573980 | NA | AAGGTGCTGGG/A | c.40_49del | p.Pro14SerfsX26 | Het | NA | Uncertain significance | 1/15,790 (0) | Neurology and Neurodevelopmental disorders* |
| chr10:92463828 | NA | AT/ A | c.2664del | p.Leu889X | Het | NA | Not present | 1/15,790 (0) | Neurology and Neurodevelopmental disorders* |
| chr10:-92556744-92561449 | NA | NA | NA | NA | Het | NA | NA | NA | Hereditary ataxia and control |
| chr10:92557209-92565408 | NA | NA | NA | NA | Het | NA | NA | NA | Syndromic congenital heart disease and control |

On the contrary, we did not identify any CNV spanning the IDE gene in the 438,250 UKBB participants. In line with these findings, only 6 small structural variants and 2 large inversions are present in gnomAD (https://gnomad.broadinstitute.org/, n=21,694 alleles).

### IDE LoF Variants in Genomics England Database

In the 15.790 neurological patients analysed in the 100,000 Genomes Project (Genomics England) screened for *IDE* rare loss of function variants (MAF < 0.01), we report 44 patients carrying heterozygous LoF variants in *IDE* (**Table2**). Among these 41 patients carried rs533083105 (p.Leu5X) and one patient carried the rs749353444 (p.Met394ValfsX29), both of them detected also in healthy controls. Two other cases inherited the frameshift variant, p.Leu889X and p.Pro14SerfsX26 from the healthy parents, mother and father, respectively.

## DISCUSSION

In this study we tested the hypothesis that *Ide* CNV (rs30920120) reported in C57B6/6J in the colonies of Jackson Laboratory and associated to a proportional *Ide* overexpression ^1^ may explain the variability of the PcomA patency that we reported during acute brain hypoperfusion in this mouse strain ^9^.

Surprisingly, all of the 63 BCCAS, MCAO and naïve C57BL/6J inbred mice from Charles River colonies (79%) and from Janvier (21%) displayed identical heterozygosity for the rs30920120 variant and did not present any *Ide* CNV (rs30920120) in our Taqman genotyping (**Figure 2**). This may be due to the relatively small sample size. Alternatively, it may be possible that the *Ide* CNV described in heterozygosity in 64% of C57BL/6J mice from the colonies of Jackson Laboratory ^1^ may not be present in different breeding laboratories. Thus, raising the intriguing hypothesis that beside a inter-and intrastrain genetic diversity there may be an additive inter-provider genetic diversity, significantly increasing the complexity of genotype–phenotype correlations.

Importantly, historically archived samples from the C57BL/6J colony suggest that *Ide* duplication has rapidly reached a high frequency in the Jackson laboratory colonies since 1994 ^1^. Notably, the Charles River laboratory received the C57BL/6J colonies in 1974 from NIH (https://www.criver.com/) and Janvier Laboratory is an independent family-owned business established in 1960 by Roger Janvier in France (https://janvier-labs.com/en/historical-overview/). Therefore, supporting the hypothesis that *Ide* CNV may be an independent event originated in and peculiar of Jackson Laboratory C57BL/6J colonies.

In our cohort of 63 C57BL/6J mice, we found a *Ide* CNV in a BCCAS mouse that during acute brain ischemia developed 2 very prominent PcomAs (**Figure 3**). However, *Ide* CNV did not segregate with the PcomA-patency phenotype in the same and different C57BL/6J stroke mouse models (MCAO) (**Figure 3**). Moreover, *Ide* expression did not significantly differ in BCCAS mice with different stroke outcomes and therefore diverse PcomA endophenotypes (absent/small/prominent/very prominent PcomA or 2 prominent PcomAs) (**Figure 4-5**).

Finally, to test the hypothesis that *IDE* gain or loss of function may have influenced a possible vascular phenotype in humans, we screened a cohort of 454.040 individuals for *IDE* CNV and 15.790 neurological patients for *IDE* LoF.

Since both *IDE* CNVs and heterozygous *IDE* LoF mutations in our cohort have been detected either in patients not presenting a pathogenic neurovascular phenotype or in healthy controls or have been inherited from healthy parents (**Table 2**), we exclude that *IDE* structural changes may critical influence a vascular trait in humans.

Thus, finally, implying that the phenotypic correlate of *Ide* CNV reported in C57BL/6J strain remains still unknown.

Importantly, IDE represents one of the principal proteases responsible for Aβ clearance ^33^. IDE activity in the human brain decreases with aging and in the early stages of AD pathology ^34^. Moreover, even modest overexpression of *Ide* in transgenic mice significantly prevents the formation and deposition of amyloid plaques in mice ^35^. Thus, considering that C57BL/6J mice are widely used as genetic background of at least 50 different AD mouse strains (https://www.alzforum.org/research-models/alzheimers-disease, **Table S3**), *Ide* duplication reported in the Jackson colonies may deeply influence interpretations of experimental results and conventional drug testing in mice, particularly in the AD field.

Finally, the brain collateral flow represents the most important survival factor during acute brain ischemia, determining the stroke lesion size and overall outcome and relies on the recruitment of collateral arteries and, particularly in the C57BL/6J mice, on PcomA patency, which, despite the Mendelian-like pattern of segregation within the strain ^9, 10^ and the numerous genetic studies, remains genetically not characterized.

In summary, our study shows that the genetic variability of the most widely used C57BL/6J mouse strain is the result of inter-and intra-strain differences and, remarkably, may also depend on the main laboratory animal suppliers. Moreover, *Ide* CNV does not critically influence PcomA phenotype in C57BL/6J BCCAS and MCAO mouse models and stroke in humans. Our findings should foster additional studies aimed at 1) improving the genetic quality control in inbred laboratory mice; 2) determining *Ide* CNV phenotypic correlate and 3) exploring the genetic determinants of PcomA caliber, which may provide a unique window into genetic determinants of collaterome in humans.

## Acknowledgements

We want to acknowledge the participants and investigators of UK Biobank, NeuroCure, Deutsches Zentrum für Neurodegenerative Erkrankungen (DZNE), Alexander von Humboldt Fellowship (to Celeste Sassi). This research was made possible through access to data in the National Genomic Research Library, which is managed by Genomics England Limited (a wholly owned company of the Department of Health and Social Care). The National Genomic Research Library holds data provided by patients and collected by the NHS as part of their care and data collected as part of their participation in research. The National Genomic Research Library is funded by the National Institute for Health Research and NHS England. The Wellcome Trust, Cancer Research UK and the Medical Research Council have also funded research infrastructure.

## Notes

### Competing Interest Statement

The authors have declared no competing interest.

