## Supplementary Tables for "*Ide* copy number variant does not influence lesion size and mortality in two C57BL/6J mouse models of cerebrovascular ischemia nor in human cerebrovascular disease. An exploratory study"

| Mouse model | PcomA phenotype | Allele X (rs30920120 C) | Allele Y (rs30920120 G) |
|---|---|---|---|
| MCAO | left small PcomA | 13, 15 | 9,2 |
| MCAO | left non-patent PcomA | 13, 25 | 10, 35 |
| MCAO | left non-patent PcomA | 12, 23 | 9, 86 |
| MCAO | left prominent PcomA | 12, 71 | 10, 39 |
| MCAO | left prominent PcomA | 14, 3 | 10, 87 |
| MCAO | left prominent PcomA | 12, 82 | 10, 58 |
| MCAO | left small PcomA | 14, 4 | 11, 63 |
| MCAO | left small PcomA | 13, 79 | 11, 15 |
| MCAO | left prominent PcomA | 14, 07 | 11, 29 |
| MCAO | left prominent PcomA | 11, 36 | 9, 52 |
| MCAO | left prominent PcomA | 13, 04 | 10, 24 |
| MCAO | left very prominent PcomA | 12, 63 | 10, 52 |
| MCAO | left small PcomA | 13, 62 | 11, 12 |
| BCCAS | 2 non-patent PcomAs | 13, 38 | 10, 72 |
| BCCAS | no PcomAs | 13, 94 | 11, 55 |
| BCCAS | 1 prominent PcomA | 14, 81 | 11, 31 |
| BCCAS | 1 small PcomAs | 14, 09 | 11, 35 |
| BCCAS | no PcomAs | 13, 32 | 11, 39 |
| BCCAS | no PcomAs | 16, 05 | 11, 47 |
| BCCAS | no PcomAs | 16, 80 | 11, 62 |
| BCCAS | prominent PcomA | 12, 52 | 10, 04 |
| BCCAS | no PcomAs | 11, 95 | 10, 46 |
| BCCAS | 1 prominent PcomA | 13, 44 | 10, 71 |
| BCCAS | 1 prominent, 1 small PcomAs | 14, 18 | 11, 23 |
| BCCAS | 1 prominent PcomA | 14, 59 | 11, 79 |
| BCCAS | 1 prominent PcomA | 13, 72 | 11, 36 |
| BCCAS | no PcomAs | 14, 20 | 11, 96 |
| BCCAS | 1 prominent PcomA | 14, 43 | 11, 64 |
| BCCAS | 2 prominent PcomAs | 13, 93 | 12, 04 |
| BCCAS | 1 prominent, 1 non-patent PcomAs | 13, 61 | 11, 49 |
| BCCAS | 2 very prominent PcomAs | 14, 27 | 11, 63 |
| BCCAS | no PcomAs | 13, 74 | 11, 43 |
| BCCAS | 1 prominent PcomA | 12, 31 | 10, 94 |
| BCCAS | 1 prominent PcomA | 13, 42 | 10, 85 |
| BCCAS | 1 prominent, 1 small PcomAs | 14, 05 | 11, 93 |
| BCCAS | 1 prominent, 1 non-patent PcomAs | 15, 31 | 12, 36 |
| BCCAS | 1 prominent, 1 small PcomAs | 15, 64 | 12, 53 |
| BCCAS | 2 prominent PcomAs | 15, 92 | 12, 27 |
| BCCAS | 1 prominent PcomA | 16, 74 | 11, 84 |
| BCCAS | 2 non-patent PcomAs | 16, 04 | 12, 39 |
| BCCAS | 2 non-patent PcomAs | 15, 24 | 12, 58 |
| BCCAS | no PcomAs | 14, 82 | 11, 27 |
| BCCAS | no PcomAs | 15, 67 | 12, 63 |
| BCCAS | no PcomAs | 16, 04 | 12, 61 |
| BCCAS | no PcomAs | 13, 40 | 10, 43 |
| BCCAS | 2 prominent PcomAs | 15, 05 | 11, 75 |
| BCCAS | 1 small PcomA | 16, 33 | 11, 00 |
| BCCAS | no PcomAs | 17, 11 | 12, 10 |
| BCCAS | 1 prominent, 1 small PcomAs | 15, 30 | 11, 59 |
| BCCAS | 1 very prominent, 1 prominent PcomAs | 13, 40 | 11, 26 |
| BCCAS | no PcomAs | 14, 07 | 11, 85 |
| BCCAS | 2 prominent PcomAs | 16, 37 | 13, 41 |
| BCCAS | 2 non-patent PcomAs | 16, 44 | 12, 79 |
| BCCAS | 1 very prominent, 1 prominent PcomAs | 14, 89 | 11, 40 |
| BCCAS | no PcomAs | 20, 07 | 13, 15 |
| BCCAS | 1 non-patent PcomA | 20, 24 | 12, 72 |
| BCCAS | 2 prominent PcomAs | 13, 97 | 10, 02 |
| BCCAS | 1 prominent, 1 non-patent PcomAs | 13, 95 | 10, 46 |
| NAIVE | no PcomAs | 13, 06 | 10, 98 |
| NAIVE | no PcomAs | 14, 18 | 11, 44 |
| NAIVE | no PcomAs | 14, 18 | 11, 16 |
| NAIVE | no PcomAs | 13, 72 | 11, 42 |
| NAIVE | no PcomAs | 13, 17 | 10, 78 |
| CONTROL | CONTROL | 0, 37 | 2, 13 |
| CONTROL | CONTROL | 0, 30 | 1, 76 |
| CONTROL | CONTROL | 0, 31 | 1, 77 |
| CONTROL | CONTROL | 0, 31 | 1, 82 |
| CONTROL | CONTROL | 0, 30 | 1, 74 |
| CONTROL | CONTROL | 0, 27 | 1, 58 |
| CONTROL | CONTROL | 0, 29 | 1, 67 |
| CONTROL | CONTROL | 0, 30 | 1, 71 |
| CONTROL | CONTROL | 0, 28 | 1, 66 |
| CONTROL | CONTROL | 0, 29 | 1, 63 |
| CONTROL | CONTROL | 0, 29 | 1, 66 |

**Table S1.** Taqman genotyping rs30920120 (C/G) results in C57BL/6J mice. MCAO, middle cerebral artery occlusion; BCCAS, bilateral common carotid artery stenosis; PcomA, posterior communicating artery

| Mouse | PcomA phenotype | number | Average CBF (Cortex and Striatum) L 0d | Average CBF (Cortex and Striatum) R 0d | Average CBF (Cortex and Striatum) L 1d | Average CBF (Cortex and Striatum) R 1d | Average CBF (Cortex and Striatum) L 1d | Average CBF (Cortex and Striatum) R 1d |
|---|---|---|---|---|---|---|---|---|
| BCCAS | 2 non-patent PcomAs | 2 | 137,11 | 135,11 | 25,36 | 12,48 | 0 | 0 |
| BCCAS | 1 prom PcomA | 9 | 116,78 | 117,27 | 33,47 | 22,30 | 37,05 | 33,16 |
| BCCAS | 1 prom PcomA | 3 | 126,73 | 122,03 | 51,27 | 22,18 | 37,94 | 49,13 |

**Table S2. Cerebral blood flow (CBF, ml/min/100g) results in different BCCAS mice at different time points.** Prom, prominent; d, day.

| AD strain name | Background | Plaque development |
|---|---|---|
| 3xTg | C7BL/6;129X1/SvJ;129S1/Sv | Plaques at 6 months |
| 5xFAD (B6SJL) | C57BL/6 x SJL | Plaques at 3 months |
| 5xFAD (C57BL6) | C57BL6 | Plaques at 3 months |
| A7 APP transgenic | C57BL/6J | Plaques at 6 months |
| ADanPP | C57BL/6J | Plaques at 3 months |
| AD-BXD | C57BL/6J X BXD | Plaques at 6 months |
| APP23 | C57BL/6 | Plaques at 6 months |
| APP23 x PS1-R278I | C57BL/6J | Plaques at 6 months |
| APP751SL/PS1 KI | 129SV x C57BL/6 | Plaques at 3 months |
| APP NL-F Knock-in | C57BL/6 | Plaques at 6 months |
| APP NL-G-F Knock-in | C57BL/6 | Plaques at 3 months |
| AppNL-G-F/MAPT double knock-in | C57BL/6J | Plaques at 2 months |
| APPPS1 | C57BL/6J | Plaques at 3 months |
| APP/PS1/rTg21221 | B6.C3 x B6.129 x FVB | Plaques at 9 months |
| AppSAA Knock-in | C57BL/6J | Plaques at 3 months |
| APPsw/0; Pdgfrβ+/- | APPsw mice on C57BL/6; Pdgfrβ+/- mice on 129S1/SvlmJ | Plaques at 9 months |
| APPSwDI x NOS2 Knock-out | C57BL/6J; C57BL/6N | Plaques at 1 year |
| APPSwe (line C3-3) | C3H/HeJ x C57BL/6J; backcrossed to C57BL/6J | Plaques at 18 months |
| APPSwe/PSEN1(A246E) | C57BL/6J x C3H/HeJ | Plaques at 9 months |
| APPSwe/PSEN1dE9 (C3-3 x S-9) | C57BL/6J | Plaques at 6 months |
| APPswe/PSEN1dE9 | C57BL/6J | Plaques at 3 months |
| APPswe/PSEN1dE9 (line 85) | C57BL/6;C3H | Plaques at 6 months |
| APP(V717I) | Originally generated on FVB/N background; available at reMYND as C57BL/6xFVB/N | Plaques at 9 months |
| APP(V717I) x PS1(A246E) | Originally generated on FVB/N background; available at reMYND as C57BL/6xFVB/N | Plaques at 3 months |
| Arc48 (APPSw/Ind/Arc) | C57BL/6 | Plaques at 2 months |
| ARTE10 | Co-injection of transgenes into B6CBF1 oocytes, back-crossed to C57BL/6 | Plaques at 3 months |
| BACE1 cKO (Hu, Yan) X 5xFAD | C57BL/6J | Plaques at 3 months |
| BRI-Aβ42 (BRI2-Aβ42) | B6C3, backcrossed to C57BL/6J | Plaques at 3 months |
| E2FAD | C57BL/6 | Plaques at 3 months |
| E4FAD | C57BL/6 | Plaques at 4 months |
| J20 (PDGF-APPSw,Ind) | C57BL/6 | Plaques at 5 months |
| mThy1-hAPP751 (TASD41) | C57BL/6 x DBA | Plaques at 3 months |
| PDAPP(line109) | C57B6 x DBA2 | Plaques at 6 months |
| PS2APP (PS2(N141I) x APPswe) | C57BL/6, DLB/2, crossed to C57BL/6 | Plaques at 9 months |
| PS/APP | B6/D2/Swe/SJL mixed background | Plaques at 6 months |
| TAS10 (thy1-APPswe) | Transgene injected into C57BL/6 x C3H oocytes, some backcrossing to C57BL/6 | Plaques at 6 months |
| TASTPM (TAS10 x TPM) | TAS10 transgene originally injected into C57BL/6 x C3H oocytes, with some backcrossing to C57BL/6 | Plaques at 6 months |
| TauPS2APP | C57BL/6, DBA/2; backcrossed to C57BL/6 | Plaques at 4 months |
| Tg2576 | B6;SJL Mixed Background | Plaques at 10 months |
| Tg2576/Tau(P301L) (APPSwe-Tau) | C57BL/6, DBA/2, SJL, SW Mixed Background | Plaques at 9 months |
| tg-APPSwe | C57BL/6J | Plaques at 12 months |
| Tg-ArcSwe | C57BL/6J | Plaques at 6 months |
| Tg-SwDI (APP-Swedish,Dutch,Iowa) | C57BL/6 | Plaques at 3 months |
| TREM2-BAC X 5xFAD | TREM2-BAC: FVB/NJ; 5xFAD: C57BL/6 X SJL | Plaques at 6 months |
| TREM2, humanized (common variant) X 5XFAD | C57BL/6 X CBA, back-crossed for at least 4 generations to C57BL/6 | Plaques at 8 months |
| TREM2, humanized (R47H) X 5XFAD | C57BL/6 X CBA, back-crossed for at least 4 generations to C57BL/6 | Plaques at 8 months |
| Trem2 KO (Colonna) x 5XFAD | C57BL/6Trem2 KO (KOMP) x APPPS1 Genetic Background: C57BL/6 | Plaques at 4 months |
| Trem2 R47H KI (Lamb/Landreth) X APPPS1-21 | C57BL6/J | Plaques at 3 months |

**Table S3. Different Alzheimer's disease (AD) C57BL6/J mouse models and respective neuropathological phenotype.**
